# Finding the neural correlates of collaboration using a three-person fMRI hyperscanning paradigm

**DOI:** 10.1101/782870

**Authors:** Hua Xie, Amber Howell, Meredith Schreier, Kristen E. Sheau, Mai K. Manchanda, Rafi Ayub, Gary Glover, Malte Jung, Allan L. Reiss, Manish Saggar

## Abstract

Humans have an extraordinary ability to interact and cooperate with others, which plays a pivotal role in societies at large. Despite its potential social and evolutionary significance, research on finding the neural correlates of collaboration has been limited partly due to restrictions on simultaneous neuroimaging of more than one participant (a.k.a. hyperscanning). A series of works now exists that used dyadic fMRI hyperscanning to examine the interaction between two participants. However, to our knowledge, no study to date has aimed at revealing the neural correlates of social interactions using a 3-person (or triadic) fMRI hyperscanning paradigm. Here, for the first time, we simultaneously measured the blood-oxygenation-level-dependent (BOLD) signal of triads (m=12 triads; n=36 participants), while they engaged in a joint drawing task based on the social game of Pictionary^®^. General linear model (GLM) analysis revealed increased activation in the brain regions previously linked with the theory of mind (ToM) during the collaborative phase compared to the independent phase of the task. Furthermore, using intersubject brain synchronization (IBS) analysis, we revealed increased synchrony of the right temporo-parietal junction (R TPJ) during the collaborative phase. The increased synchrony in the R TPJ was observed to be positively associated with the overall team performance on the task. In sum, our novel paradigm revealed a vital role of the R TPJ among other ToM regions during a triadic collaborative drawing task.

## 1. Introduction

“Teamwork makes the dream work,” coined originally by John Maxwell (Maxwell, n.d.), emphasizes the importance of teamwork and collaboration as opposed to working independently or competing with others to achieve shared goals. As one of the defining factors of human beings (Dunbar, 2016), collaboration is pivotal for societies at large to bring industrial and scientific advancement (Dodgson, 1994; Lewis et al., 2012). A better understanding of the neural basis for how we collaborate could not only help us advance our knowledge of social cognition, but also provide us ways to enhance collaboration and teamwork.

Almost all social cognitive neuroimaging studies have utilized single-person experimental paradigms, where a set of pre-recorded and time-locked events is used to examine the neural underpinnings of a particular social behavior across participants (e.g., Rilling et al., 2002; Walter et al., 2004). Such single-person experimental paradigms have significantly enhanced our understanding of several aspects of social cognition, such as social reasoning, social reinforcement, and social emotion (for review, see Adolphs, 2003). Several neuroimaging studies have already highlighted the importance of the theory of mind (ToM) network and the mirror neuron system (MNS) for inferring social intentions and joint action (Van Overwalle and Baetens, 2009). The ToM network, consisting of the precuneus, temporo-parietal junction (TPJ) and medial prefrontal cortex (mPFC), is thought to be responsible for understanding others’ internal mental states (Arioli et al., 2018). On the other hand, the MNS, consisting of areas such as the inferior frontal gyrus (IFG), premotor cortex (PMC), inferior parietal lobule (IPL) and superior parietal lobule (SPL), is considered essential to understanding others’ overt actions and their underlying causes by internal neural simulations (Alcalá-López et al., 2018).

The single-person neuroimaging experiments, however, are inherently limited in examining the full essence of social cognition as back-and-forth interactions, and continuous adaptation in everyday social interactions are entirely absent in a single-person paradigm (Hari and Kujala, 2009). A natural solution to examining the intricate dance of social interactions is hyperscanning, i.e., simultaneous recording of brain activities from multiple participants during real-time social interactions (Montague et al., 2002). Since then, a number of studies have started to adopt hyperscanning paradigm together with inter-subject brain synchrony analysis (IBS; brain-to-brain coupling) to study social interactions during collaboration. Higher IBS is commonly observed in positive coordinated interpersonal behavior, such as cooperation (Cui, Bryant, & Reiss, 2012; Miller et al., 2019; Saito et al., 2010; Toppi et al., 2016), and is linked to greater shared understanding and intentionality (Nguyen et al., 2019; Tang et al., 2015). Although hyperscanning is gaining popularity, two issues still remain. The first issue is that a large portion of previous hyperscanning studies (and *all* previous fMRI-based hyperscanning studies) have focused on the interaction between a pair of participants, or a *dyad*. Without getting into the ongoing debate on whether dyad actually constitutes a group or not (Moreland, 2010; Williams, 2010), we ought to acknowledge that dyadic interactions, albeit useful, are limited in studying more complex phenomenon such as social rejection or mediation. Thus, transitioning from a dyad to a *triad* fundamentally shape the way individuals think and act (Simmel, 1908), and triads are more suitable for exploring these more complex in-group interactions (Moreland, 2010). A handful of EEG and NIRS hyperscanning studies have gone beyond dyads. For example, Jiang et al. (2015) and Nozawa et al. (2016) studied leader emergence in a triad, and Dikker et al. (2017) examined EEG brain synchronization among a group of 12 students during class. However, the potentially richer information available from fMRI hyperscanning has not been tapped to date to evaluate the neural correlates of social collaboration among three (or more) individuals. The second issue pertains to the ecological validity of experiments. As with most neuroimaging studies, well-controlled lab experiments provide invaluable information about specific brain circuits, but recently more naturalistic experiments with high ecological (or real-life) validity (Hari and Kujala, 2009) are gaining favor.

To fill some of these gaps, here we conducted a 3-person fMRI hyperscanning experiment where participants played a version of the word-guessing improvisation game of Pictionary (Angel, R. & Everson, 1985) and took turns to collaborate and draw a given set of action words (or verbs). We have previously used this social improvisation game to study neural correlates of spontaneous improvisation and figural creativity (Saggar et al., 2017, 2015). Here, we modified the paradigm to include three phases: independent phase, where participants worked on their own to draw each given word; evaluation phase, where they could see and evaluate other team members’ drawings from the independent phase; and lastly collaboration phase, where participants worked jointly on a shared screen and took turns to re-draw the word in a collaborative manner (Figure 1). Importantly, in the collaboration phase, participants could see each other’s drawings in real time, thereby allowing us to examine brain-to-brain coupling in real time. We also included control words (e.g., draw a spiral) in the independent phase to contrast for basic visuospatial activity. After the experiment, participants rated other team members (including themselves) on performance. Further, to estimate task performance, the drawings generated by participants (both during independent and collaborative phase) were rated on the ease of guessing and originality by an independent panel of judges (authors R.A. and H.X.).

**Figure 1.**
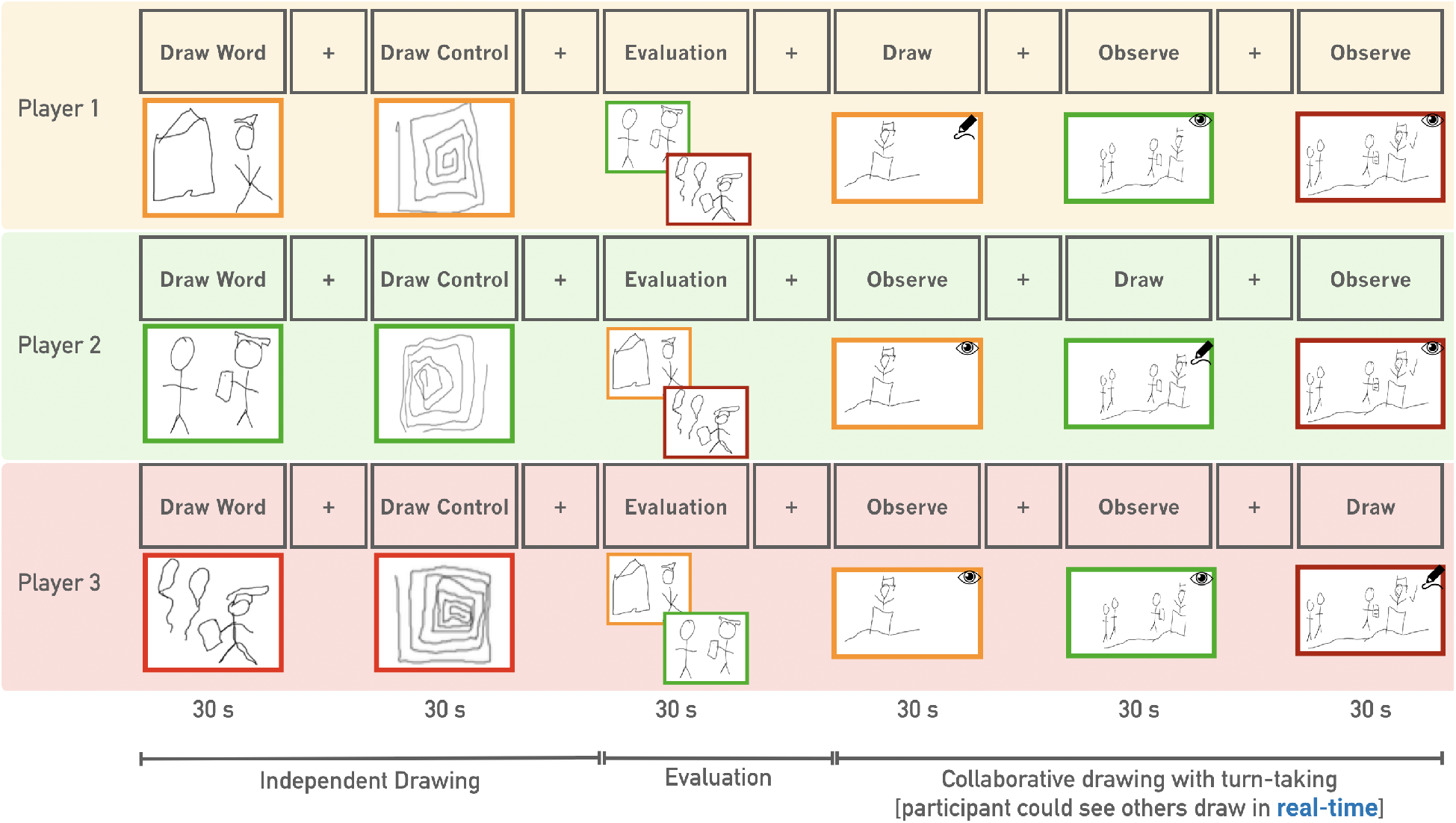
An illustration of the multiplayer joint improvisation paradigm for one verb (graduate). Participants used an MR-safe tablet and stylus to complete the drawing. A total of nine verbs were drawn in 3 runs. For each verb, there were three phases: independent drawing, evaluation and collaborative drawing. In the collaboration phase, participants took turns to draw (order was counterbalanced), while everyone could see each other’s drawings in real time.

Like spoken and written language, drawing (or sketching) is a form of communication and externalizing ideas (Tversky, 2002). During the collaboration phase, participants shared their understanding by drawing, which might have further enabled them and others to discover hidden relations and generate novel insights (Masaki Suwa et al., 2000). Hence, a vital objective of this study was to uncover the neural underpinnings of collaboration by contrasting independent drawing with collaborative drawing. To achieve this goal, we first performed a GLM analysis to identify the brain networks associated with collaborative drawing as compared to independent drawing. We then conducted an IBS analysis to reveal brain regions that show increased synchrony during the collaboration phase. We hypothesized that the social collaboration would recruit regions in the ToM network and elicit higher IBS among teammates with better coordination and better team performance.

## 2. Results

Here we conducted a 3-person fMRI hyperscanning experiment where participants played a version of the word-guessing improvisation game of Pictionary (Angel, R. & Everson, 1985) and took turns to collaborate and draw a given set of action words (or verbs). Participants first worked independently to draw each given word, followed by evaluating other players’ drawings, and finally took turn to re-draw the word in a collaborative manner (Figure 1). Importantly, in the collaboration phase, participants could see each other’s drawings in real time. In this section, we first provide results from a GLM analysis that was done to identify the brain regions associated with collaborative drawing as compared to independent drawing. We then reported results from the IBS analysis, which reveals brain regions that show increased synchrony during the collaboration phase.

### 2.1. Behavioral data

Thirty-six right-handed adults (20M, 16F; 27.44±4.98y) were recruited for our study and written informed consent was obtained from all participants. Behavioral assessments were conducted to assess the participants’ general intelligence, personality, positive/negative affect and figural creativity, and post-scan questionnaires were collected (e.g., rating performance of all team members). See Methods and Supplementary Tables (S1-6) for detailed demographic and behavioral assessment information. The drawings generated during the fMRI session were evaluated by two raters on the scales of originality (uniqueness across all participants) and usefulness (i.e., level of ease for another person to guess the word represented by the drawing). Furthermore, a composite team performance score was created by multiplying the two scores of originality and usefulness. Supplementary Table S7 provides information on range and distribution of these scores across triads.

### 2.2. Examining neural correlates of collaboration using GLM analysis

We first set out to identify regions associated with collaboration while drawing. To control for basic visuospatial processing, we first contrasted independent and collaboration drawing with control drawing (i.e., drawing spiral shapes). Two contrasts were created: independent contrast (i.e., independent – control drawing) and collaborative contrast (i.e., collaborative – control drawing). We then examined the difference between these two contrasts using GLM. Results are presented in Figure 2. We conducted an additional GLM analysis to directly compare collaborative with independent drawing (i.e., without using control drawing) and similar results were observed (as shown in Supplementary Figure S1).

**Figure 2.**
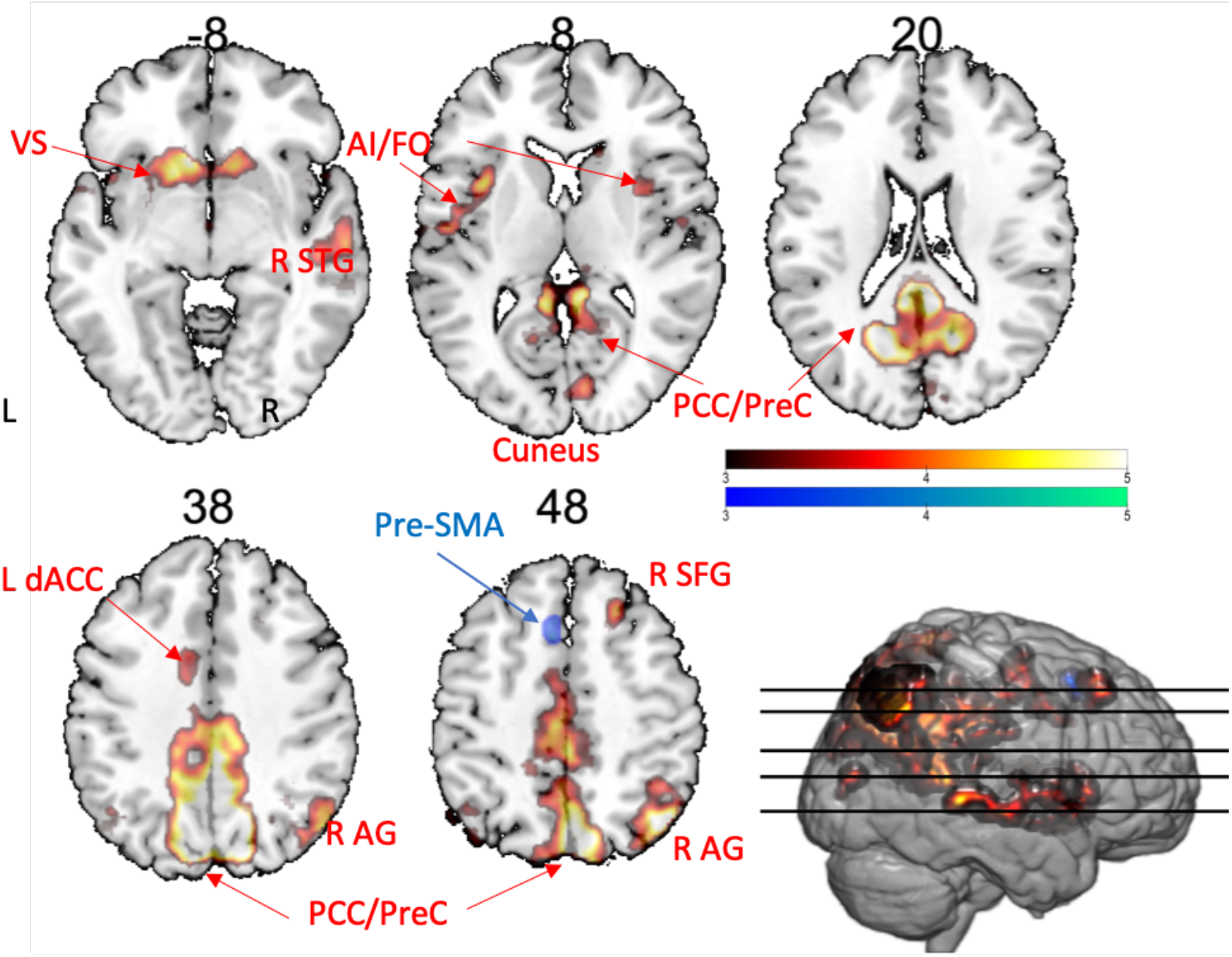
Neural correlates of collaborative versus independent drawing. The red-yellow scale depicts contrast of collaborative > independent drawing, while the blue-green scale represents the reverse contrast.

As shown in Figure 2 and Table 1, collaboration > independent contrast revealed increased activity in eight clusters including the posterior cingulate cortex/precuneus (PCC/PreC), right angular gyrus (R AG), bilateral ventral striatum (VS), right superior temporal gyrus/middle temporal gyrus (R STG/MTG), bilateral anterior insular/frontal opercular cortex (AI/FO), left dorsal anterior cingulate cortex (L dACC), right superior frontal gyrus (R SFG) and right cuneus. For the reverse contrast, i.e., independent > collaboration, increased activity in the pre-supplementary motor area (pre-SMA) was revealed.

**Table 1.** Summary of contrast pattern of collaborative improvisation versus independent drawing (and reverse). The MNI coordinates with peak z-statistic and number of voxels within each cluster are also reported.

| Collaboration > Independent | Brain region | Z-MAX | X (mm) | Y (mm) | Z (mm) | Cluster size (number of voxels) |
| --- | --- | --- | --- | --- | --- | --- |
| 1 | PCC/PreC | 6.39 | 13.5 | -60.5 | 27.5 | 10,630 |
| 2 | R AG | 5.79 | 45.5 | -66.5 | 49.5 | 1,096 |
| 3 | VS | 5.17 | -8.5 | 21.5 | -6.5 | 1,045 |
| 4 | R pSTG/MTG | 4.62 | 67.5 | -34.5 | -2.5 | 789 |
| 5 | L AI/FO | 4.63 | -34.5 | 11.5 | 9.5 | 576 |
| 6 | L sLOC/AG | 4.11 | -48.5 | -72.5 | 41.5 | 205 |
| 7 | R SFG | 4.31 | 23.5 | 27.5 | 51.5 | 143 |
| 8 | R Cuneus | 4.18 | 11.5 | -82.5 | 9.5 | 115 |
| Independent ><br>Collaboration |  |  |  |  |  |  |
| 1 | Pre-SMA | 4.4 | -4.5 | 15.5 | 53.5 | 107 |

**Table 2.** Summary of correlation analysis between team-level IBS and composite performance scores. MNI coordinates of center of gravity (COG) for each ROI and associated *p*-value.

| Seed region | COG X (mm) | COG Y (mm) | COG Z (mm) | $p$-value |
| --- | --- | --- | --- | --- |
| R sLOC/AG | 50.33 | -60.06 | 36.14 | 0.0073 |
| R TPJ | 51.54 | -56.59 | 15.78 | 0.039 |

### 2.3. Examining neural correlates of collaboration using IBS analysis

To further investigate the neural coupling among teammates, we applied inter-subject brain synchronization (IBS) analysis to identify regions that were synchronized across teammates during the collaborative phase. After parcellating the brain using the Shen atlas (Shen et al., 2013), the mean region-wise timeseries were z-scored and IBS was estimated using the pairwise Pearson’s correlation coefficients. To find the brain regions that were significantly synchronized across participants within each triad, we performed permutation testing to estimate difference between IBS estimated from true triads with those estimated from “fake” triads (created by shuffling team labels). Positive False Discovery Rate (FDR) correction was performed to correct for multiple hypothesis testing (Storey, 2002), and ROIs with a FDR-corrected Q-value smaller than 0.05 are shown in Figure 3.

**Figure 3.**
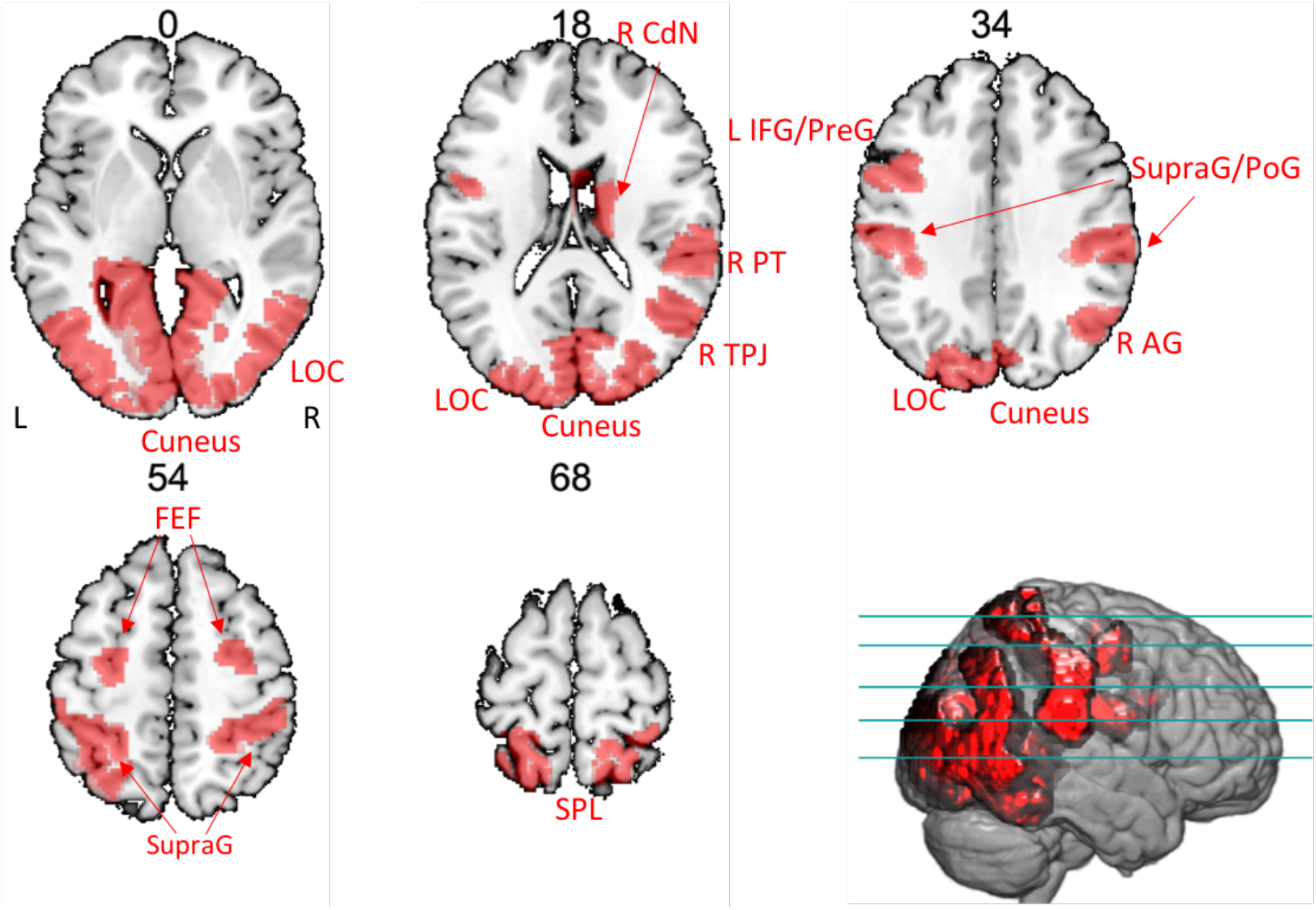
A binary mask of regions with significantly higher within-triads IBS during the collaboration phase (Q < 0.05) are shown in red. These regions included lingual gyrus (LG), occipital pole (OP), cuneus, occipital fusiform gyrus (OFG), lateral occipital cortex (LOC), supramarginal/postcentral gyrus (SupraG/PoG), right planum temporale (R PT), right temporoparietal junction (R TPJ), right angular gyrus (R AG), frontal eye field (FEF), and left inferior frontal gyrus/precentral gyrus (L IFG/PreG).

To test if the observed within-triad IBS was primarily driven by the residual task structure shared within triad across triads, we also performed IBS analysis on the timeseries derived from the independent phase. No significant results were obtained for any brain region during the independent phase.

### 2.4. Examining relation between increased IBS and team performance

To examine the brain-behavior relationship we calculated correlation between the behavioral team performance and the degree of observed IBS across participants within each triad. We hypothesized that the teams with higher IBS during the collaboration phase would perform better on the task. For the team performance, we collected two metrics. First metric was team-level composite performance score of the sketch, as defined by the product of usefulness and originality scores of each triad’s final drawing from the collaboration phase. These sketches were rated by two judges on the scales of usefulness and originality as per Saggar et al. (2015), with an inter-rater reliability > 0.9 (using intraclass correlation coefficient; ICC). The second metric was pairwise rating from a post-scan questionnaire, where each participant rated the performance of their team members during the entire paradigm.

We computed the team-averaged IBS and correlated each team’s mean IBS with team-level composite performance score averaged across all verbs. Figure 4 shows two ROIs with significant correlation with composite performance scores, i.e., right angular gyrus/superior lateral occipital cortex (R AG/sLOC, rho = 0.75, *p* = 0.007) and right temporo-parietal junction (R TPJ, rho = 0.63, *p* = 0.039). Additionally, we observed a statistically nonsignificant relation between IBS of the R TPJ and pairwise performance rating (rho = 0.30, *p* = 0.092), as shown in supplemental Figure S3.

**Figure 4.**
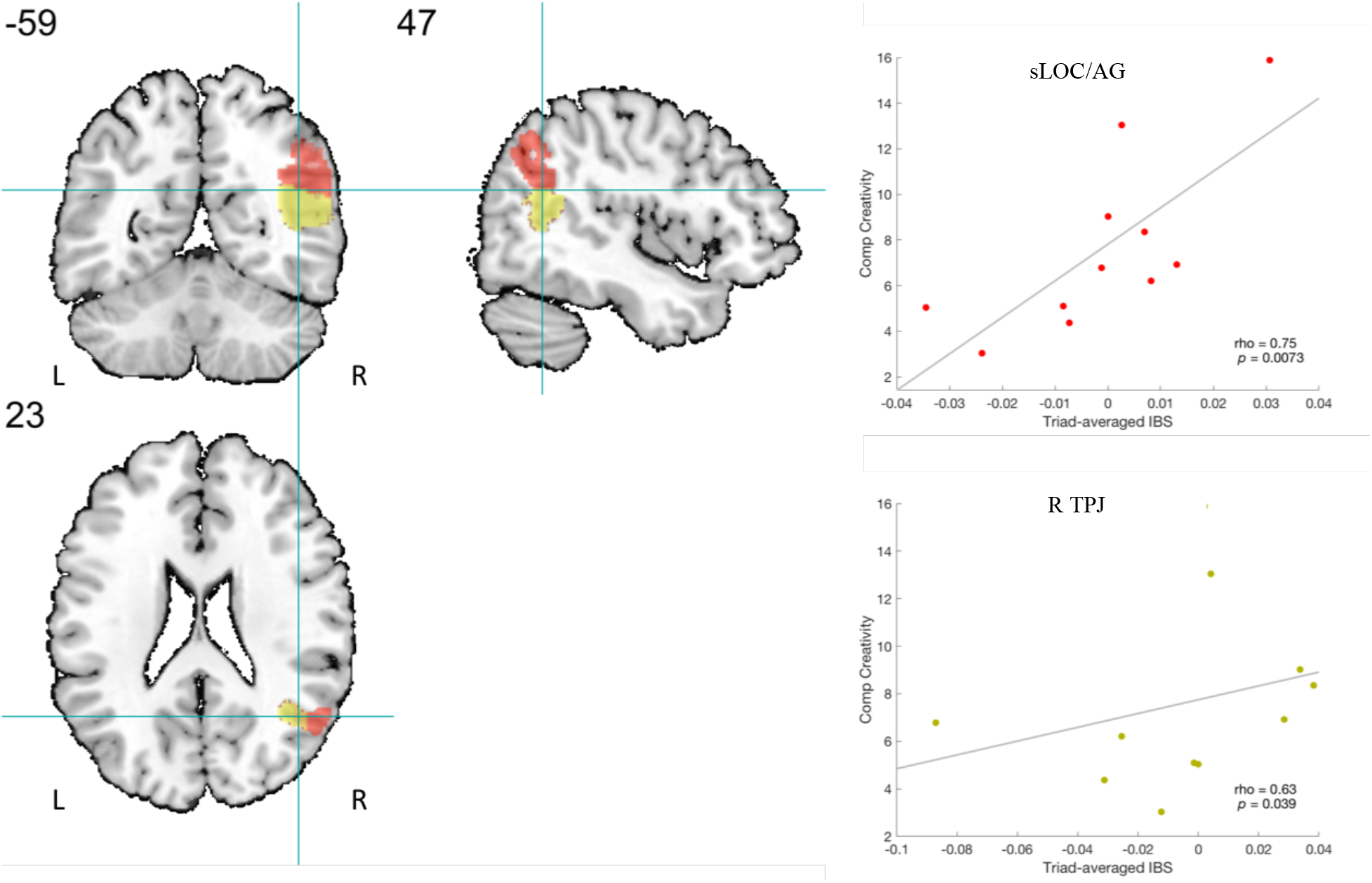
Triad-averaged IBS vs. averaged composite creativity score. Red: R sLOC/AG (rho = 0.75, *p* = 0.007, uncorrected); Yellow: R TPJ (rho = 0.63, *p* = 0.039, uncorrected), *p*-value remained significant after removing the leftmost point.

## 3. Discussion

It is a well-known phenomenon that the presence of others drastically changes how we behave, e.g., the bystander effect (Hortensius and de Gelder, 2014) and the joint Simon effect (Dolk et al., 2014). Yet, the majority of social neuroscience is limited to single-person neuroimaging experiments, which in turn prevents us from studying more realistic complex social phenomena (Todorov et al., 2006). Here, for the first time, we performed a 3-person fMRI hyperscanning experiment to study the neural correlates of collaboration using a social game of Pictionary. Our results highlighted the importance of the ToM network, especially the right temporo-parietal junction (R TPJ) during the 3-person social interaction.

Using single-subject neuroimaging experiments, neuroscientists have traditionally focused on studying our ability to understand the mental states of others as an observer. What is lesser known is how our brains dynamically adapt when we are actively engaged in a social interaction. This lack of understanding about inter-brain interaction has also been referred to as the “dark matter” of social neuroscience (Schilbach et al., 2013). Recent development of hyperscanning tries to address the issue by simultaneously scanning multiple participants during real-time social interactions. To quantify between-subject brain coupling, inter-subject brain synchronization (IBS) is typically used (Finn et al., 2018; Hasson et al., 2012; Salmi et al., 2013). As a model-free approach, IBS does not require a priori experimental design and is well suited to study natural and open-ended social interactions (Nummenmaa et al., 2018). Here, using IBS in a 3-person hyperscanning paradigm, we examined the neural correlates of collaboration while participants were engaged in a social improvisation paradigm of Pictionary.

The theory of mind (ToM) network has been known to play a central role in social cognition. Two core regions in the ToM network include the medial prefrontal cortex (mPFC) and bilateral TPJ, while the precuneus, inferior frontal gyrus, and anterior temporal lobes are also considered part of the ToM network (Schurz et al., 2014). Although part of the same network, the two core regions (mPFC and TPJ) have been thought to assume distinct social cognitive functions (Babiloni and Astolfi, 2014; Chauvigné et al., 2018; Zheng et al., 2018). It has been hypothesized that while the TPJ is linked with assessing transient mental inferences about other people’s goals/beliefs, the mPFC is responsible for assessing trait judgments of others and self (instead of immediate actions/goals; Van Overwalle, 2009). Further, among the bilateral TPJs, the right TPJ (R TPJ) has been specifically shown to play a critical role in establishing social context and is of particular interest given its robust activation across a broad range of social cognitive tasks (Schurz et al., 2014).

Our GLM results confirmed the role of ToM network during the collaboration phase, as we saw increased activations in the PCC/Precuneus and R angular gyrus (R AG) as compared to the independent phase. We also observed increased activations in the regions of AI/FO, dorsal ACC (dACC), and ventral striatum (VS). It should be noted that the AI/FO, ACC, and the amygdala are considered to be part of the emotional salience network, suggesting the involvement of subjective emotion such as happiness, empathy, and uncertainty (Craig, 2009). Moreover, stronger activation in the VS was previously found during live interaction between participants and experimenters (Redcay et al., 2010). To our surprise, we did not observe increased activation in the bilateral mPFC and TPJ during collaboration. The absence of the mPFC seemed to coincide with an earlier study by Schippers et al. (2009), who also reported negative evidence of involvement of mPFC during a gestural communication task. This is likely because the mPFC is involved in inferences of enduring characteristics (personality traits) rather than transient information (actions/goals), as suggested by Van Overwalle (2009). Despite the absence of increased activation in the TPJ, we did observe increased synchronization of R TPJ during collaboration. Further, the increased R TPJ synchronization among teammates was later observed to be positively associated with team performance.

Our IBS analysis revealed a distinct set of brain regions being involved during collaboration as compared to activation patterns observed using GLM analysis. Besides the putative stimulus-locked IBS of primary visual cortex and somatosensory cortex, the dorsal attention network (DAN) including superior parietal lobule (SPL) and frontal-eye-field (FEF) was also observed to be more synchronized during collaboration. Increased IBS in the DAN potentially indicates a top-down control of visual attention, given the shared understanding of the current word to be drawn and similar expectations based on the pre-existing drawing (Corbetta and Shulman, 2002). Higher synchronization of right TPJ and left IFG was also observed, both of which are part of the ToM network. In addition to its role in the ToM network, R TPJ is also postulated to be associated with the ability to reorient attention to unexpected stimuli (Corbetta et al., 2008). Recent neuromodulation studies have concluded that the R TPJ could play an overarching role in both domains of attention as well as social cognition (Krall et al., 2016). Thus, it is possible that higher IBS in the R TPJ during collaboration could be partially attributed to the emergence of an unexpected/salient idea put forth by other teammates. However, we also observed the degree of synchronization in the R TPJ was positively related to team’s overall performance. The observed relation between brain-to-brain coupling of the R TPJ among teammates and their performance points towards the likelihood that in our study the R TPJ may indeed be related to better coordination within the team. Our conclusion is also supported by the finding by Tang et al. (2015), where higher IBS in R TPJ is linked to greater shared intentionality between individuals.

It is intriguing that little spatial overlap was found comparing GLM activation pattern with IBS co-fluctuation pattern. However, such lack of overlap was partially intentional; as we removed task-related signals before conducting IBS analysis. This was done to reduce the influence of shared baseline conditions on IBS, otherwise the IBS pattern would mainly reflect activation pattern itself (Pajula et al., 2012). Such procedures have been previously adopted so that IBS pattern could reveal additional information beyond just activation patterns (Saito et al., 2010).

### Limitations & future work

As is always the case, the ecological validity and relatively unconstrained interactions between participants come with a cost. First, even though turn taking is a common phenomenon in our daily interactions, such as conversation (Sacks et al., 1978), it poses some difficulties on the behavioral and neuroimaging data analysis, especially when our design involved three individuals without fixed roles such as drawer and observer (Nozawa et al., 2016). For example, individuals’ response strategy could depend on the drawing order. Although we counterbalanced the drawing order across all participants during the collaboration phase, the order of drawing could still affect participants’ strategy and engagement level, and consequently influenced IBS and GLM outcomes. For example, during the collaboration phase, we sometimes observed participants repeat their drawings from the independent phase (e.g., for the first drawer) or simply add details instead of adding a novel component (e.g., for the third drawer). Some of these strategies could be subjective (e.g., whether it is a new idea or a modification of a previous idea or adding details) and cannot be readily inferred from their drawings alone. Future work may benefit to include a questionnaire of participants’ response strategy at the end of each task block or limit the response strategy to encourage participants to always generate new ideas. Besides, concerns may arise from the fact the drawing order might also lead to difference in the engagement level. As each participant only had one chance to draw during the collaborative phase, one might be more attentive before his/her turn as compared to afterwards. Indeed, it has been shown that active viewing may result in different IBS pattern as compared to passive viewing (Nummenmaa et al., 2014). The authors found active viewing led to increased IBS in precentral gyrus, parietal cortex and DAN. Interestingly, we found increased IBS in similar regions and thus suggesting that participants in our task were likely attentive even during observation. Overall, given our engaging game-like experimental design and the counterbalancing of order of drawing across words, we anticipate the order effect should be mitigated to a large degree.

The second issue pertains to the sample size of this study, even with 36 participants we only have 12 triads. Lack of power due to relatively small number of triads could result in less robust brain-behavior correlation analysis. Thus, care should be taken when interpreting the observed relation between increased IBS in the R TPJ and overall team performance of the triads.

A promising future avenue for this work is to study the direction of information flow between participants rather using between-brain Granger-causality analysis (Schippers et al., 2010). The IBS, as measured here using correlation, only quantifies pairwise relationships and does not take full advantage of triadic interactions. Future investigations should look into methods that might better characterize the team interaction beyond pairwise relationship and understand the brain dynamics associated with such complex social interaction (e.g., using Topological Data Analysis; Saggar et al., 2018).

We also acknowledge that individual differences (e.g., personality, drawing skills, and creativity level) could play an important role in team performance, hence subsequent neuroimaging analysis. Take creativity as an example, Taggar (2001) showed that groups made up of more creative members were found to be more creative on the team level, whereas Harvey (2014) argued that team creative performance also depends on the effective integration of team members. While our analysis focused on the second perspective using team performance as a proxy of team integration/collaboration, it is likely that individual difference also affected the overall team performance (e.g., more creative or open participants might contribute more than in the collaborative phase). It would be interesting to study the influence of team composition in future study (Xue et al., 2018). Additionally, it should also be noted that due to a technical error, acquisition parameters differed across sites. We attempted to account for multi-site variability by modeling sites as a fixed effect in the higher-level GLM analysis (see Methods 4.4.2). IBS analysis should be less susceptible to such differences due to the averaging of voxel-wise timeseries within ROIs. Nonetheless, one-way ANOVA analysis revealed no significant group differences in IBS across sites (see Methods 4.4.3).

Lastly, given the nature of the task (drawing), despite our best effort of keeping participants from moving, head motion could have influenced the results. We adopted a rigorous approach for removing head movement artifacts by first using ICA-AROMA (Pruim et al., 2015) and then censoring high-motion frames (frame displacement > 0.5mm) as well as dropping high-motion subjects (more than 30% frames discarded) from the analysis.

### Conclusions

In this study, we performed the first ever 3-person fMRI hyperscanning experiment to study the neural underpinning of social collaboration using a Pictionary-like joint drawing paradigm. We found collaboration-related activation in ToM regions including PCC, precuneus and R AG. We also highlighted importance of the R TPJ, and its relationship with positive interpersonal collaboration outcome in the form of better team performance rating. In sum, triadic hyperscanning joined by open-ended task paradigm offers a unique avenue for neuroscientists to disentangle complex everyday group interactions.

## 4. Methods

### 4.1. Participants

Thirty-six healthy adults were recruited in our study (age: 27.44±4.98y, 16 female), who were randomly assigned to twelve triads. Participants were considered eligible if they were 18-45 years old, right-handed, with no history or neurologic or psychiatric illness. The participants were not trained in any forms of visual arts. The written consent forms were obtained as approved by Stanford University’s Institutional Review Board (Human Subjects Division).

### 4.2. Multi-subject hyperscanning paradigm

The collaborative verb-drawing fMRI task is a multi-player version of Pictionary^®^ game developed based on a previous study by Saggar et al. (2015). The goal of the task is to draw a verb independently and collectively for others to guess. Nine verbs were drawn over three runs (3 verbs per run), i.e., run1: snore, graduate, accelerate; run2: whisper, salute, vote; run3: redial, boil, pinpoint. Each verb can be further broken down into two phases: independent phase (3 conditions) and collaborative phase (2 conditions), each phase consisting three 30s-blocks. Task blocks were separated by a fixation period jittering around 7-8s.

Independent phase: in the independent phase, participants performed the task independently, and were unable to see the each other’s screen. The three conditions within independent phase are independent drawing, control drawing, and inspiration, with each block lasting for 30s. For the independent drawing condition, the participants were instructed to draw a verb (e.g. graduate) within 30s, and the verb was shown on the top-left corner of the screen and present throughout the block. Participants were instructed to fully utilize the given 30s-block and continue to add elements to the illustration instead of finishing early. It was followed by a 30s control drawing block, during which participants drew spiral shapes (e.g., spiral triangle, square, circle). The control block was designed to control for basic motor and visuospatial aspect of the independent drawing as well as to balance the cognitive demands, as the subjects were instructed to draw spiral shapes without intersecting any lines. In the inspiration condition, participants were asked to view the drawings of the other two participants and mark the interesting features.

Collaboration phase: in the collaboration phase, participants performed the task collaboratively by sharing a screen through Internet. Collaboration phase consisted of three 30s-blocks. During each block one participant drew the same verb and other two observed, with the goal to collaboratively create a single drawing. The order of drawing was pseudorandomized across verbs so that every subject had an equal opportunity to draw first.

An MR-safe tablet was used for drawing, with which participants practiced prior to the scan. The tablet was designed specifically for a previous study by Saggar et al. (2015) with a KEYTEC 4-wire resistive touch glass connected to a Teensy 2.0 with custom firmware, which was connected to the console through a USB port. We recorded every stroke and associated timestamp which we used to reconstruct drawings. Participants were told to hold the tablet with their left hand, and stylus in right hand. Padding was provided under the participant’s upper arms and under the tablet to provide support while subjects were holding the tablet and to restrict additional movement of hands.

The triad was scanned simultaneously on three 3T GE scanners using 32-channel head coils (Discovery MR750, General Electric, Milwaukee, WI) at two different locations. Two subjects were scanned at Stanford University’s Richard M. Lucas Center and the third one at the Stanford Center for Cognitive and Neurobiological Imaging. The T2*-weighted functional images were collected with repetition time (TR) = 2s, echo time (TE) = 30ms, flip angle (FA) =77°, field of view (FOV) = 23.2 x 23.2 cm, 40 axial slices with slice thickness of 2.9mm. Due to a technical error, the acquisition parameters were slightly different across scanners (see Table S6 for details). Further analysis was carried out to ensure the difference in acquisition parameters did not confound our GLM (section 4.4.2) or IBS results (section 4.4.3). High-resolution T1-weighted structural images were acquired with TE = 3.2ms, TR = 8.2ms, FA = 15°, 1mm isotropic voxel size, FOV = 25.6 x 25.6 cm, 186 axial slices with slice thickness of 1mm.

For stimuli presentation, synchronization and communication between participants, a web-server application was developed using JavaScript (node.js; https://nodejs.org/en/) and was run using the Hypertext Transfer Protocol (http). For establishing communication protocols between participants and the server, we used JavaScript based Socket.IO libraries (https://socket.io/). For drawing, the Canvas application programming interface (API) was used. Canvas elements can be used for real-time animation, game graphics, data visualization, etc. (for more details, please see https://developer.mozilla.org). We plan to release the entire code on our lab’s GitHub page (https://github.com/braindynamicslab) upon publication.

### 4.3. Behavioral assessments

#### 4.3.1. General Intelligence

The Wechsler Abbreviated Scale of Intelligence-II (WASI-II; Wechsler, 2011) is used to measure general intelligence. Only two subtests, vocabulary and matrix reasoning, were administered.

#### 4.3.2. Personality

The NEO Five-Factor Inventory-3 (NEO-FFI-3; McCrae & Costa, 2004) contains 60 item and measures of the five domains of personality (neuroticism, extraversion, openness, agreeableness, and conscientiousness).

#### 4.3.3. Affect

The Positive and Negative Affect Schedule (PANAS; John & Julie, 2004) is a self-report questionnaire that consists of two 10-item scales to measure both positive and negative affect.

#### 4.3.4. Figural creativity

The figural form of the Torrance Tests of Creative Thinking (TTCT-F; Torrance, 1974) is a paper-pencil based test used to assess creativity including figural fluency, flexibility, originality, and elaboration, and an average standard score can be computed as an overall score for figural creativity.

#### 4.3.5. Pairwise/triadic behavioral metrics

The pairwise performance rating between the participants was collected in a post-scan questionnaire, in which we asked them to rate all subjects’ performance on a five-point scale (1-5), with 1 being very strongly dissatisfied and 5 being strongly satisfied. As for the triadic performance, two raters evaluated the final drawings from the collaboration phase and generated two additional behavioral ratings: originality (the number of the unique elements in the drawing) and usefulness (the level of ease for another person to guess the word represented by the drawing). The drawings were rated on a five-point scale as 1 means not original/useful, and 5 means very original/useful.

### 4.4. fMRI data analysis

#### 4.4.1. Preprocessing

We used fMRIPrep v1.1.4 (https://fmriprep.readthedocs.io/en/stable/index.html; Esteban et al., 2019), a preprocessing toolbox built on a combination of existing neuroimaging software, to preprocess our MRI data. Briefly, a reference volume of functional data and its skull-stripped version were generated using a custom methodology of fMRIPrep. Six rigid-body head motion parameters with respect to the functional reference image were estimated using mcflirt (FSL; Smith et al., 2004), followed by slice-timing correction using 3dTshift from AFNI (Cox RW, 1996). The functional reference was then co-registered to the structural reference using flirt (FSL; Smith et al., 2004), with nine degrees of freedom to account for any remained distortions in the functional reference. Functional data were then normalized to MNI152 space (2 mm isotropic) following the spatial smoothing with Gaussian kernel (FWHM = 6mm). ICA-AROMA, a robust technique to automatically remove motion artifacts using independent component analysis (Pruim et al., 2015) and tested on both task-based and resting-state data, was applied through a “non-aggressive” way.

#### 4.4.2. General linear modeling (GLM)

For the general linear modeling, we used FEAT (FMRI Expert Analysis Tool; version 6.00) to identify the brain activations associated with each condition. First four volumes were discarded, and the remaining were high-pass filtered (128s, 0.008Hz). For the subject-level analysis, boxcar model (one for each of the five conditions) and canonical hemodynamic response function (HRF) was used for GLM estimation with temporal derivative enabled for any potential lag of HRF. Since our task involved drawing, despite our effort to keep subjects still, it was inevitable that motion had some correlation with the task condition (e.g., motion during drawing condition being higher than observation condition). A prior study has showed that even moderate task-correlated motion has a deleterious effect on true activations, if motion parameters are included as covariates (Johnstone et al., 2006). Hence, we decided to use data preprocessed by ICA-AROMA given its robust performance reducing motion artifacts (Parkes et al., 2018), and also included physiological signal (average white matte, CSF signal, and the derivatives) as the nuisance regressors in our GLM analysis. To further reduce the effect of residual head motion artifacts, we censored the volumes based on the frame displacement (FD) and discarded any data points with FD greater than 0.5mm, as deleting high-motion frame has shown to improve the quality of GLM estimation (Siegel et al., 2014). Among all 108 datasets that we planned to collect (36 participants x 3runs/participant), seven runs were excluded from further analysis either due to subjects dropping out or missing behavioral data. Another four datasets were dropped due to excessive motion (%volume discarded > 30%). All participants had at least one run entering GLM analysis as well as IBS analysis. Group-level z-statistic images were thresholded at Z = 3.1 and familywise error correction (FWE) corrected threshold of *p* = 0.05. To control for a possible scanner site effect, we included the scanner site as a nuisance covariate in the group-level analysis, and no significant clusters were found to be associated with site effects in the main contrasts (FWE corrected *p* > 0.05).

#### 4.4.3. Inter-subject brain synchronization (IBS)

We parcellated the functional data using the Shen atlas (Shen et al., 2013) to generate ROI timeseries. More specifically, we first generated a binary group mask from all the datasets and overlaid the group mask on the Shen atlas. We removed ROIs outside the field of view and ROIs that had little overlap with our group mask (#overlapped voxels < 10). A total number of 241 ROIs remained. Physiological noise, task structure regression, and bandpass filter (0.008 - 0.09Hz) were implemented in one single step using 3dTproject (AFNI). Frame censoring was performed by replacing high-motion frames (FD > 0.5mm) by nans. Then, we z-scored the residual timeseries and concatenated the collaborative periods across all available runs in order to compute inter-subject brain synchronization (IBS). Here, IBS was quantified as the Pearson’s correlation coefficients between timeseries of a given ROI between two subjects (Hasson et al., 2008), which captures the time-locked neural co-fluctuation across subjects. IBS were computed between all subject pairs, including IBS between subject pairs from the same triad (within-triad IBS) as well as IBS between subject pairs from different triads (between-triad IBS) and the difference between the two based on the true triad labels (IBS_true_). We ran 50,000 permutation tests to generate the null distribution of IBS by shuffling the triad labels, recomputed the IBS based on the shuffled labels (IBS_shuffled_), and the *p*-value was quantified as the proportion of IBS_true_ greater than IBS_shuffled_. The *p*-values were subsequently corrected for multiple comparisons via positive FDR (Storey, 2002) using Matlab function mafdr for all ROI pairs, and Q-value of 0.05 was used as the threshold which yielded a total number of 42 significant IBS as shown in Figure 3. Exact same procedures were taken while using the time points from the independent phase to test if IBS was driven by the residual task structure shared within triad or any timing differences across triads. No IBS survived after positive FDR correction. Additionally, we employed the one-way ANOVA to test if any combinations of scan parameters across different sites had significant effect on within-triad IBS strength, and we did not find any IBS pairs survived multiple comparison correction (Q > 0.05).

We were also interested in exploring if the IBS of those 42 ROIs can be associated with any pairwise or triad-level task performance. For that purpose, we first averaged the performance rating between a pair, and subtracted the triadic mean to mitigate any potential triad effect. To reduce the effect of motion on pairwise IBS, we defined the pairwise FD which is calculated as the inner product of FD timeseries between subject pairs during the collaboration phase. The pairwise FD was regressed out from the IBS, and residual IBS was then demeaned within the triad and correlated against demeaned pairwise rating Spearman’s rank correlation.

We further tested if the triad-level performance metrics could be related to the residual triad-averaged IBS. Here, we defined an averaged composite creativity score as the product of originality and usefulness scores averaged across all words. Given the limited number of triads (11 triads as one triad was reduced to a dyad when discarding the high-motion runs), we utilized the robust correlation toolbox (Pernet et al., 2013) to compute the two-tailed Spearman’s correlation between triadic composite creativity score and triad-averaged IBS values and significance was determined by 1,000 bootstrapping.

## Acknowledgements

Funding by K99/R00 to M.S., Stanford MediaX to A.L.R. and a gift from the Albert Yu and Mary Bechmann Foundation.

## Author Contribution

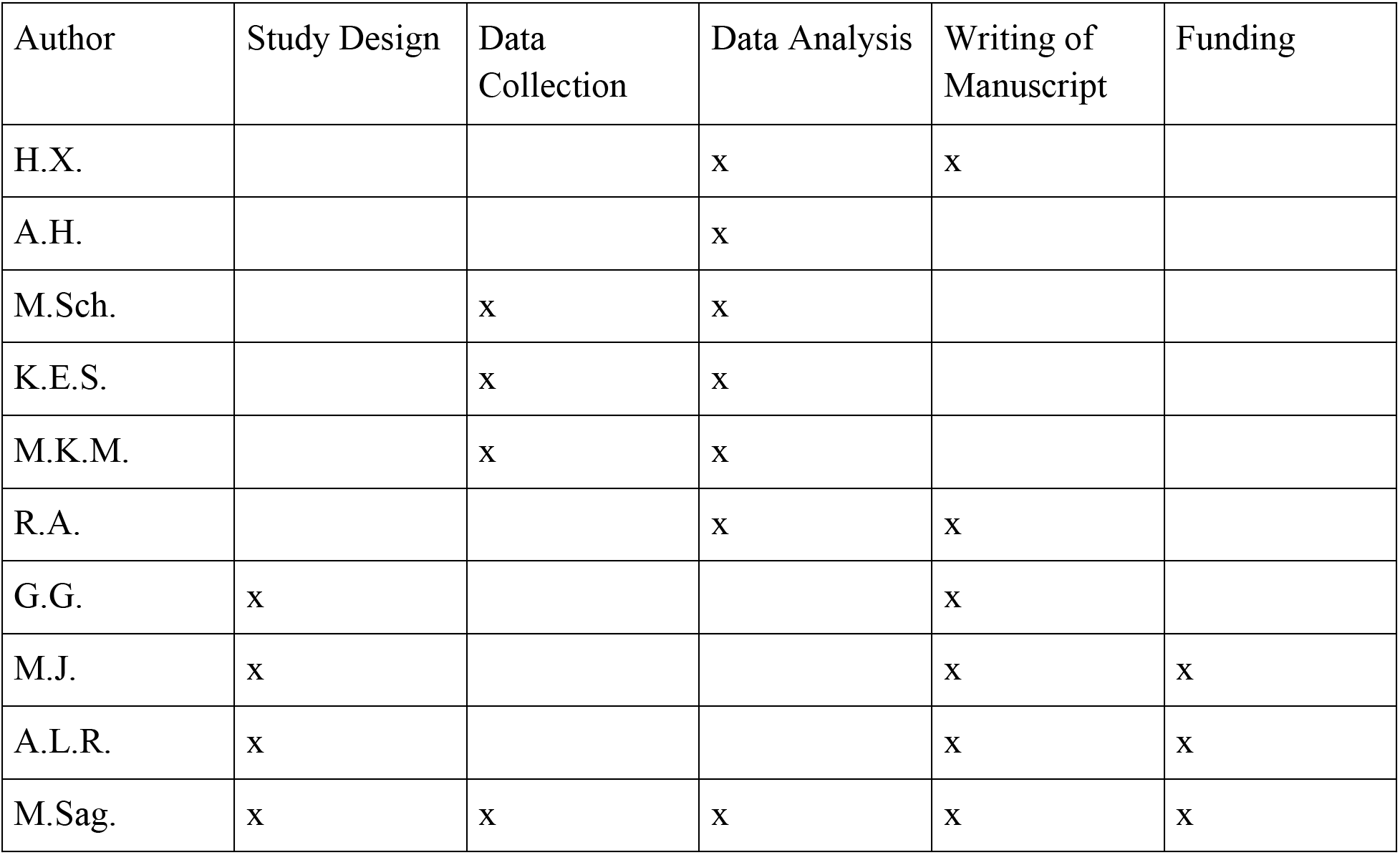

